# SPPUSM: An MS/MS spectra merging strategy for improved low-input and single-cell proteome identification

**DOI:** 10.1101/2023.06.29.547039

**Authors:** Yongle Chen, Zhuokun Du, Hongxian Zhao, Wei Fang, Tong Liu, Yangjun Zhang, Wanjun Zhang, Weijie Qin

## Abstract

Single and rare cell analysis provides unique insights into the investigation of biological processes and disease progress by resolving the cellular heterogeneity that is masked by bulk measurements. Although many efforts have been made, the techniques used to measure the proteome in trace amounts of samples or in single cells still lag behind those for DNA and RNA due to the inherent non-amplifiable nature of proteins and the sensitivity limitation of current mass spectrometry. Here, we report an MS/MS spectra merging strategy termed SPPUSM (same precursor-produced unidentified spectra merging) for improved low-input and single-cell proteome data analysis. In this method, all the unidentified MS/MS spectra from multiple test files are first extracted. Then, the corresponding MS/MS spectra produced by the same precursor ion from different files are matched according to their precursor mass and retention time (RT) and are merged into one new spectrum. The newly merged spectra with more fragment ions are next searched against the database to increase the MS/MS spectra identification and proteome coverage. Further improvement can be achieved by increasing the number of test files and spectra to be merged. Up to 18.2% improvement in protein identification was achieved for 1 ng HeLa peptides by SPPUSM. Reliability evaluation by the “entrapment database” strategy using merged spectra from human and *E. coli* revealed a marginal error rate for the proposed method. For application in single cell proteome (SCP) study, identification enhancement of 28%-61% was achieved for proteins for different SCP data. Furthermore, a lower abundance was found for the SPPUSM-identified peptides, indicating its potential for more sensitive low sample input and SCP studies.

## Introduction

Organs, tissues, and cultured cells all consist of a variety of cells with distinct molecular and functional properties^1^. Characterizing cellular heterogeneity by single-cell technology is key for understanding the normal physiology and disease pathology of living organisms^2–5^. Recent discoveries in neuroscience, developmental biology, and stem cell biology by single-cell genomic and transcriptomic studies have urged scientists to expand single-cell investigations to a broader set of “omics”^6–10^. Single-cell proteomics (SCP) is aimed at uncovering the protein heterogeneity present among cells that would have otherwise been masked in bulk samples^11–13^. However, due to the non-amplifiable nature of proteins and the limited sensitivity of the currently used proteome strategy^14^, SCP still faces challenges in achieving in-depth sequencing coverage and high throughput. Many attempts have been made to boost the SCP sequencing depth, including reducing overall protein loss in sample preparation caused by surface adsorption^3,14–23^, improving ion transfer efficiency^24^ and enhancing detection sensitivity of the mass spectrometry (MS) system^24–29^. In addition to the above “wet experiment techniques”, data processing is another important part of promoting proteome identification in SCP^30,31,32^. Match between runs (MBR) algorithm in MaxQuant has been shown to improve the SCP identification scale^33^, although further improvement in false discovery rate (FDR) control is needed^34^. Spectral clustering is also capable of increasing protein identification by grouping similar MS/MS spectra and transferring the peptide identity among different runs^35–37^. However, both MBR and spectral clustering require successful MS2 level identification in at least one “run” for the propagation of the peptide sequence among replicate runs. Therefore, they are not applicable to peptides without identified MS/MS spectra throughout the entire study.

Most proteomic search engines^38–41^ that are extensively applied now were initially established for bulk sample data, which intrinsically require high-quality spectra produced by sufficient precursor ions. However, owing to the extremely low amount of peptides and limited ions transferred into the MS analyzer in SCP, the quality of MS/MS spectra is usually lower than that of bulk samples^32^. The obviously reduced number of fragment ions in the MS/MS spectra consequently results in low SCP identification rates^42^. Some researchers have focused on more efficient accumulation of precursor ions, for example, by prolonging the maximum ion injection time of MS1 aquistion^25,42^ to improve MS/MS fragmentation. However, this strategy is followed at the cost of reducing the MS scan speed and losing acquired spectra, which may jeopardize the peptide quantification accuracy. Payne et al. found that the fragmentation patterns of peptides in SCP differ from those of the bulk sample^32^. In bulk samples, the MS/MS spectra of the same peptide tend to display relatively high reproducibility. Similar fragment ions and intensity distributions can be obtained among replicate runs. However, for SCP, fewer annotated ions and poor signal-to-noise ratios are usually observed in the spectra, which may result in lower similarities among spectra from replicate runs, especially for the low-quality ones^32^.

Based on the above discovery, we proposed a same precursor-produced unidentified spectra-merging (SPPUSM) strategy for more efficiently exploiting the MS/MS spectra obtained in replicate samples and to achieve deeper proteome identification for low-input and single-cell studies. MS/MS spectra merging was previously used as a part of the de novo sequencing workflow, especially when multifragmentation modes were applied, since enriched fragment ions were needed for peptide identification without a database^43–46^. In our strategy, all the unidentified MS/MS spectra obtained from replicate MS runs were extracted and filtered by precursor mass, retention time (RT) and charge to find possible MS/MS spectra produced by the same precursor ions. Next, the spectra of the same precursor ion were merged into one new spectrum with more abundant fragment ions and higher intensities by exploiting the complementary nature of different spectra generated in replicate tests (Scheme 1). After re-performing the database searching, a maximum of 43 and 95% new PSM can be successfully identified to support up to 18% and 61% enhancements in protein identification for 1 ng HeLa digested peptides and single HeLa cells. Unlike spectra clustering and MBR strategies that require identified spectra for peptide sequence propagation, SPPUSM focuses on elucidating completely unidentified MS/MS spectra to improve peptide sequencing. Since our strategy simply involves post-run data processing of the obtained spectra, and because dozens to hundreds of replicates are usually conducted in SCP studies, we expect the SPPUSM strategy to be used to improve SCP spectra interpretation and enlarge the SCP identification scale without the need of extra “wet” experiments.

**Scheme 1.**
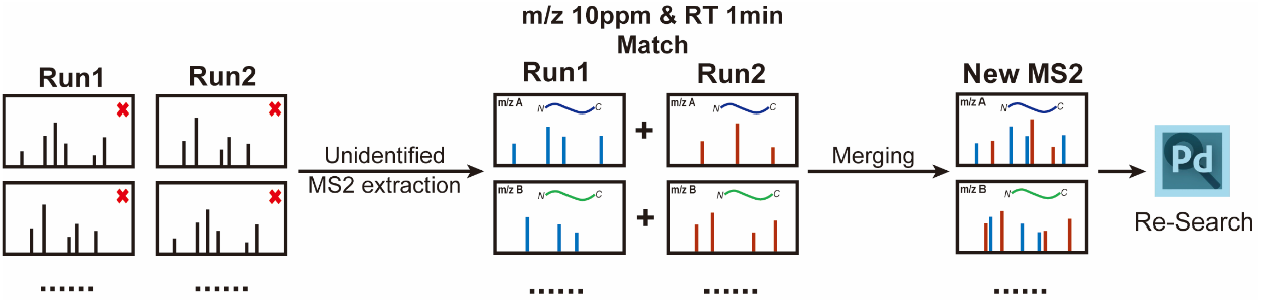
Schematic overview of the SPPUSM workflow.

## Materials and methods

### Cell culture and cell sorting

HeLa cells and HEK293T cells were separately cultured in DMEM (Gibco, UK) supplemented with 10% fetal bovine serum (FBS, Gibco, UK) and 1% penicillin-streptomycin (HyClone, USA). The cells were cultured in a humidified incubator (Thermo Fisher Scientific, USA) with 5% CO_2_ at 37 °C. For bulk samples of HeLa cell digestion, protein extraction and digestion was performed by following the FASP protocol^47^. For the single-cell analysis of HeLa cells, before adding the single cells, 0.2 mL polymerase chain reaction (PCR) tubes (Corning, USA) were pretreated by overnight incubation at 37°C with a 10 µL mixture of 5 ng/µL dodecyl-beta-D-maltoside (DDM, J&K Scientific, China) and 1 ng/µL trypsin (Promega, USA) in water to saturate the inner surface and to prevent sample loss by nonspecific adsorption. After 8 µL of the solution was removed, a BD FACSAria II cell sorter (BD biosciences, USA) was employed to directly isolate single cells into the PCR tubes and the tubes were kept at -80 °C overnight for cell lysis. The samples were next defrosted on ice, incubated at 72 °C for 30 min, and centrifuged at 6000 g at 4 °C for 5 min^48^. Next, 0.4 µL of 10 ng/µL trypsin in water was added to the solution and incubated overnight at 37 °C. Lastly, 2 µL of 0.5% formic acid (Sigma Aldrich, USA) in water (v/v) was added to the solution to quench the reaction and adjust its pH for ESI-MS analysis.

### LC-MS/MS analysis

For 500-0.2 ng HeLa digest as well as synthetic standard peptides (Synpeptide CO., LTD, China), LC-MS analysis was performed using an Orbitrap Q-Exactive HF mass spectrometer (Thermo Fisher Scientific, USA) coupled with an EASY-nLC 1000 system (Thermo Fisher Scientific, USA). A housemade 15 cm-long LC column of i.d. 150 µm packed with 1.9 µm C18 packing particles (Dr. Maisch GmbH, Germany) was used for peptide separation. For single-cell analysis, LC-MS analysis was performed using an Orbitrap Exploris 480 mass spectrometer (Thermo Fisher Scientific, USA) equipped with a FAIMS Pro interface (Thermo Fisher Scientific, USA) and an EASY-nLC 1000 system, FAIMS cycling between CVs of -45 V and -65 V every 1.3 s. A housemade 20 cm-long LC column of i.d. 75 µm packed with 1.9 µm C18 packing particles was used for peptide separation. For peptide elution, a 75 min gradient of 6-40% buffer B (80% acetonitrile with 0.1% formic acid) was used for both LC systems. The samples were spiked with iRT standards (Biognosys, Switzerland) for retention time calibration. MS measurements were performed in data-dependent acquisition mode. For samples from 500 ng to 10 ng of HeLa digest and synthetic standard peptides, MS1 spectra were acquired at a 120,000 resolution with a scan range of 300 to 1400 m/z, AGC target of 3e6 and maximum injection time (ITmax) of 50 ms. Precursors with an intensity of greater than 2.5e4 were selected for MS2 analysis as a ‘top30’ method, isolated at a 1.6 m/z isolation width, collected at an ITmax of 20 ms or AGC target of 5e4, then fragmented with a normalized collision energy of 27, and the resulting fragment ions were analyzed in the Orbitrap at a resolution of 15,000. For 1 ng and 0.2 ng of HeLa digest and single-cell analysis, MS1 spectra were acquired at 120,000 resolution with a scan range of 300 to 1400 m/z, AGC target of 3e6 and ITmax of 80 ms. Precursors with an intensity > 5.0e3 were selected as a ‘top25’ method, isolated at a 1.6 m/z isolation width, collected at an ITmax of 100 ms and AGC target of 5e5^25,42^, then fragmented with a normalized collision energy of 27, and the resulting fragment ions were analyzed in the Orbitrap at a resolution of 15,000.

### MS data processing and spectra merging

All the MS raw files to be merged were first processed using MaxQuant (version 2.0.2.0) for feature detection and database searching. MS/MS spectra were searched against the UniProtKB/Swiss-Prot human database (downloaded on 20220201). The first search peptide tolerance was set to 20 ppm, and the main search peptide tolerance was set to 4.5 ppm. Trypsin was specified as the enzyme that allowed up to two missed cleavages. The carbamidomethylation of cysteine was specified as a fixed modification, and the protein N-terminal acetylation and oxidation of methionine were considered variable modifications with a total of two variable modifications per peptide. The false discovery rate was set at 1% at both the peptide and protein levels. Following the search, ‘allPeptides.txt’ and ‘evidence.txt’ in the result text files were further processed. According to the “evidence.txt” file, all the information of the unidentified MS/MS spectra, including the scan number and MS1 features (precursor charge, m/z and RT), were extracted from the “allPeptides.txt” file. After the raw files were converted into mascot generic format (mgf) files with MSConvert (version 3.0.2), the unidentified MS/MS spectra in different files of replicate runs using the same sample amount were extracted and matched using the information in the “allPeptides.txt” file based on the following criteria. The RT tolerance and MS1 precursor mass tolerance were set as 1 min and 10 ppm, respectively. Next, the corresponding unidentified MS/MS spectra in different files that met the filtering criteria were merged into a new spectrum containing all the fragment ion peaks of the original spectra. For fragment ion peak merging, a 10 ppm mass tolerance was used. Specifically, if the mass difference was within 10 ppm, the two fragment ion peaks were considered the same, and their intensities were accumulated. Otherwise, both peaks were retained in the new spectra. Lastly, the new mgf files containing the merged spectra were re-searched using Proteome Discoverer (PD, version 2.4) against the same database that was previously used, since MaxQuant is not compatible with mgf file processing. Considering the limited number of merged MS/MS spectra that can be obtained from replicate tests, spectra from the corresponding original raw files were added to the mgf file of the merged spectra for database searching to facilitate FDR filtering. PD was used for protein/peptide identification comparison before and after SPPUSM processing to prevent inconsistency between PD and MaxQuant.

## Results and discussion

The idea of using merged spectra originated from the assumption that a lower spectra similarity is expected in samples of lower amount. Therefore, combining the complementary b and y ions by merging the spectra from different LC-MS runs may result in improved spectra elucidation. The spectra similarities among replicate runs were first investigated using different amount of HeLa digested peptides. The spectra similarities among replicate runs were first investigated using different amount of HeLa digested peptides. As showed in Figure S1A-C (Supporting information), one co-identified peptide from 0.2 ng, 10 ng and 250 ng samples manifested different spectra patterns. Spectra from 0.2 ng sample exhibited less fragment ions, lower S/N ratio and more importantly, higher stochastic identification of b and y ions. In contrast, higher spectra similarity and reproducibility of b and y ions was observed for 10 and 250 ng samples. This trend was more clearly displayed by cosine scores of all the peptides identified from 0.2, 10 ng and 250 ng samples in Figure S1D (Supporting information). As the sample amount decreased from 250 ng to 0.2 ng, the cosine scores dropped accordingly with a sharper decrease for 0.2 ng samples, indicating more complementary spectra merging may be obtained for trace amount samples.

Precursor mass tolerance and RT tolerance are the two key factors that influence the results of MS/MS spectra matching and merging. 10 ppm tolerance was set for precursor ion and fragment ion matching, which was the same or even more strict than that required in data dependent acquisition-based database searching for data obtained by Orbitrap analyzer^39,49^. Next, 1 ng and 0.2 ng HeLa digested peptides were used to investigate the appropriate RT tolerance setting. Identified PSMs in replicate runs were matched using the above mass tolerance and 2, 1 and 0.5 min RT tolerance to calculate correct match rates. As showed in Figure S3, nearly 99% correct matching rates were achieved for both 1 ng and 0.2 ng HeLa digested peptides when 1 and 0.5 min RT tolerance were used. Therefore,1 min RT tolerance was adopted to allow more candidate spectra. The feasibility of using the SPPUSM workflow for improving spectral quality and facilitating peptide identification was first evaluated using synthetic standard peptides. Table 1 listed the sequence, m/z and RT of the synthetic peptides. Four types of peptides were specially designed with two isoforms (the same amino acid composition but different sequences) for each type to mimic peptides that shared the same m/z and similar RTs in the real sample, which are most likely to be misidentified. After data processing by the SPPUSM workflow, as shown in Figure 1A, merging the unidentified spectra of replicate runs of the same peptides led to the identification of the corresponding peptides in the sample. The typical MS/MS spectra of the four peptides before and after merging in Figure S2 indicate the potential capacity of the SPPUSM workflow to increase annotated fragments in spectra. By contrast, merging the unidentified MS/MS spectra from MS runs of different peptides did not result in the identification of any peptides (Figure 1B), even though the peptides shared the same m/z and close RTs and therefore were most likely to lead to incorrect spectral merging when using MS1 feature matching.

**Table 1.** Synthetic standard peptides used for the evaluation of the SPPUSM workflow.

| No. | peptide sequence | mass (Da) | retention time |
| --- | --- | --- | --- |
| 1.1 | ASLINPGQEK | 1055.5611 | 21±1 min |
| 1.2 | ASGNLIPQEK | 1055.5611 | 22±1 min |
| 2.1 | GYGSNFVVGER | 1183.5622 | 49±1 min |
| 2.2 | GYGVVFSNGER | 1183.5622 | 49±1 min |
| 3.1 | MAFDLERPGVPVENR | 1728.8617 | 38±1 min |
| 3.2 | MAFRPGVVDLPEENR | 1728.8617 | 38±1 min |
| 4.1 | GYLQPYTDEEEDALIHR | 2047.9487 | 38±1 min |
| 4.2 | GYTLADEEEDPLIYQHR | 2047.9487 | 37±1 min |

**Figure 1.**
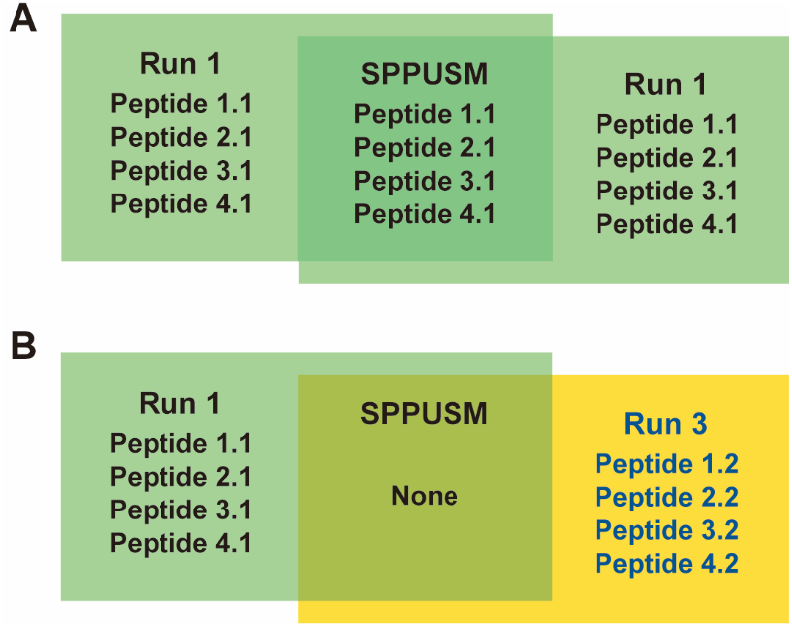
Evaluation of the SPPUSM workflow using isoforms of synthetic standard peptides. (A) SPPUSM results of merging spectra from replicates of the same peptides. (B) SPPUSM results of merging spectra from peptide isoforms.

After demonstrating the feasibility of using the SPPUSM workflow for improving spectral identification with standard peptides, the efficiency of the proposed strategy was further evaluated using more complex HeLa cell-digested peptides. After the spectral merging of the data acquired from two technical replicates by the SPPUSM workflow, 3.4% to 26.1% newly identified peptide spectrum matches (PSMs) were obtained using different amounts of digested peptides (Figure 2A). Although the number of new PSMs decreased with decreasing peptide amounts, the ratio of PSM enhancement increased and reached the top level (26.1%) at 1 ng of peptides, indicating the advantage of using the SPPUSM workflow for improving spectral identification for low sample amounts. Further decreasing the peptide amount to 0.2 ng resulted in a 6.2% drop of the PSM enhancement to 19.9%. Next, the number of files used for spectral merging was investigated. Five replicate tests were conducted for 250 to 0.2 ng HeLa peptide samples. As the number of merged files increased, the ratios of the new PSMs exhibited a clear enhancement for all the tested sample amounts except for 250 ng (Figure 2B). The ratio of new PSMs is calculated by dividing the number of new PSMs by the total number of PSMs in the original files.

**Figure 2.**
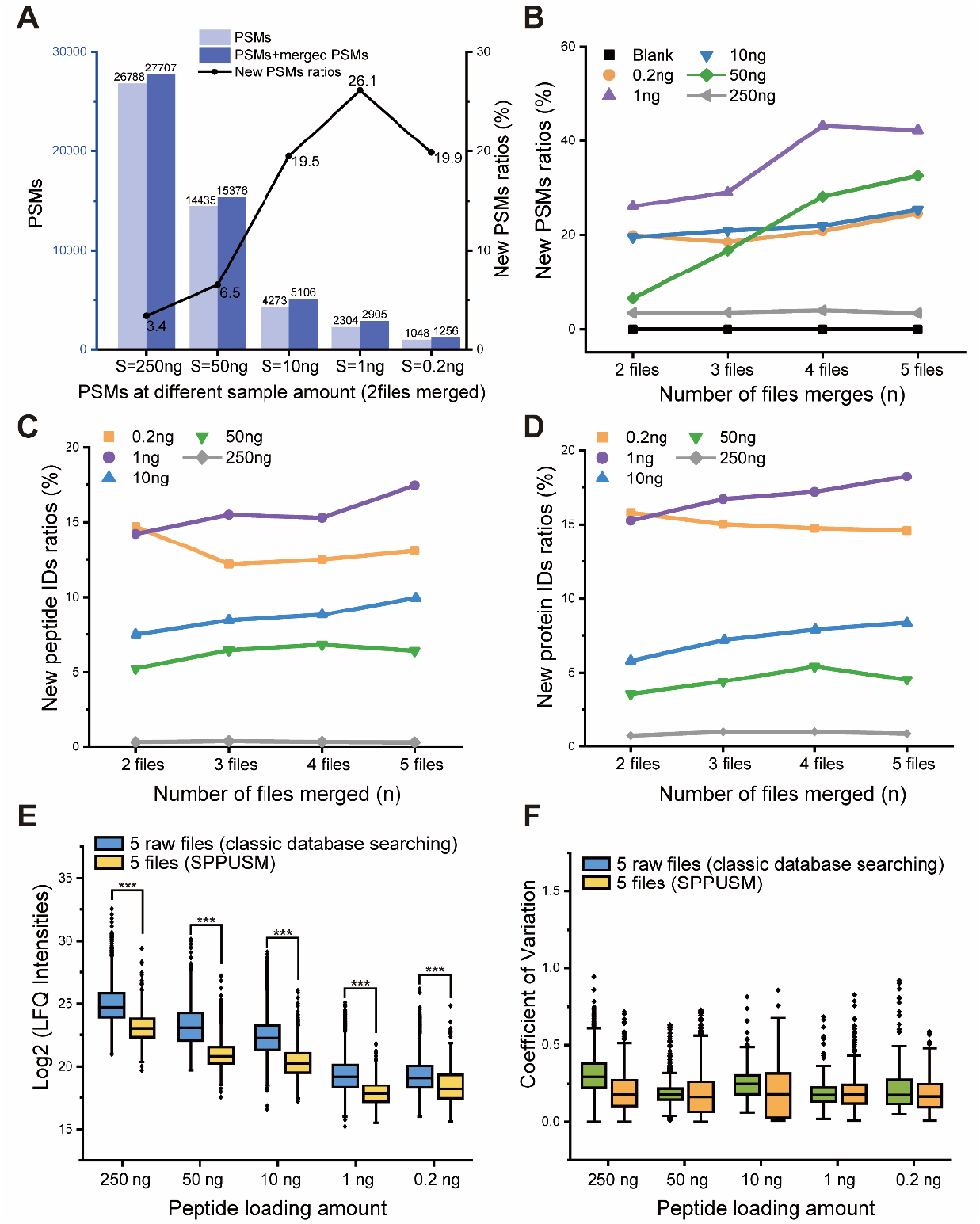
PSMs, unique peptides and protein groups identified by the classic database searching strategy and by the SPPUSM workflow using 250 ng, 50 ng, 10 ng, 1 ng and 0.2 ng digested HeLa cell peptides. (A) Number of PSMs obtained and ratio of new PSMs. (B) Percentage of new PSMs obtained by the SPPUSM workflow using different number of replicates. (C and D) Percentage of new peptides and proteins obtained by the SPPUSM workflow. (E) Distributions of log2-transformed LFQ intensities of the peptides identified by classic database searching vs. SPPUSM. The *** symbol indicates a statistically significant difference (*P* < 0.01). (F) Distributions of the CVs of peptide LFQ intensities.

For 250 ng peptides, only 3-4% new PSMs were obtained by merging two to five files, which may be attributed to the high quality of the MS/MS spectra obtained using bulk peptides; therefore, the room for improvement by spectral merging was limited. In contrast, 50 ng peptides displayed more distinct increase in PSM identification after spectral merging, given that 33% new PSMs were obtained by merging the spectra from five replicates. Further enhancement was achieved by even lower sample amounts and 1 ng peptides provided the highest improvement of 43% new PSMs. Similar rising trends were found for protein and peptide identification using the SPPUSM workflow (Figure 2C and D). Increased numbers of new proteins and peptides were identified from the merged spectra as the number of replicates rose from two to five, since more spectral merging resulted in richer fragment peaks (Figure S4) and facilitated spectral identification. The maximum enhancement of 18.2% for proteins and 17.4% for peptides was achieved at 1 ng of sample. As expected, we found that the newly identified peptides by the SPPUSM workflow indeed displayed lower intensities compared with the peptides obtained by the classic database searching method (Figure 2E), which is consistent with the lower abundance of peptides resulting in fewer fragment ions and therefore impeded their identification by the classic database searching method. After spectral merging, the number of fragment ions increased and thus facilitated the interpretation of the spectra. Further comparison of the coefficients of variation (CVs) of the intensities of the peptides obtained from replicate runs revealed that the SPPUSM workflow exhibited similar CVs as the classic database searching method in Figure 2F. Lower than 25% median CVs were achieved in replicated runs by the SPPUSM workflow, indicating the reliability of the newly identified peptides.

Theoretically, the new peptides identified by the SPPUSM workflow should be low abundant peptides that can also be identified using the classic database searching method by increasing the sample amount. Therefore, we prepared a sequence pool by combining the identified peptides from 500-50 ng samples using the classic database searching method. Figure 3 revealed that almost all of the newly identified peptides by the SPPUSM workflow can be efficiently covered by the pool. Especially for 0.2 ng samples, the coverage for SPPUSM was even 10% higher than that obtained by the classic database searching method. Further evaluation was conducted on the reliability of the SPPUSM workflow by “entrapment database” strategy using merged spectra from one HeLa cell raw file and different numbers of *E. coli* raw files that were searched against the human proteome database. Since no correct identification will be obtained by merging HeLa cell and *E. coli* spectra after excluding the homologous peptides between them, the error rate of this strategy can be estimated by counting the number of the SPPUSM peptides identified in this way and then using the following equation. A descending trend was discovered for the error rate with the increasing number of *E. coli* files used for the SPPUSM workflow (Figure S5). The lowest error rates reached 2.9% when four *E. coli* files were used for merging, indicating that relatively reliable results can be obtained.

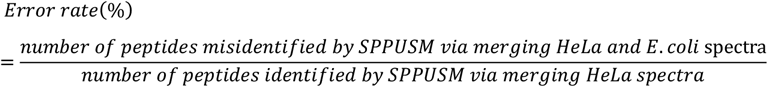

**Figure 3.**
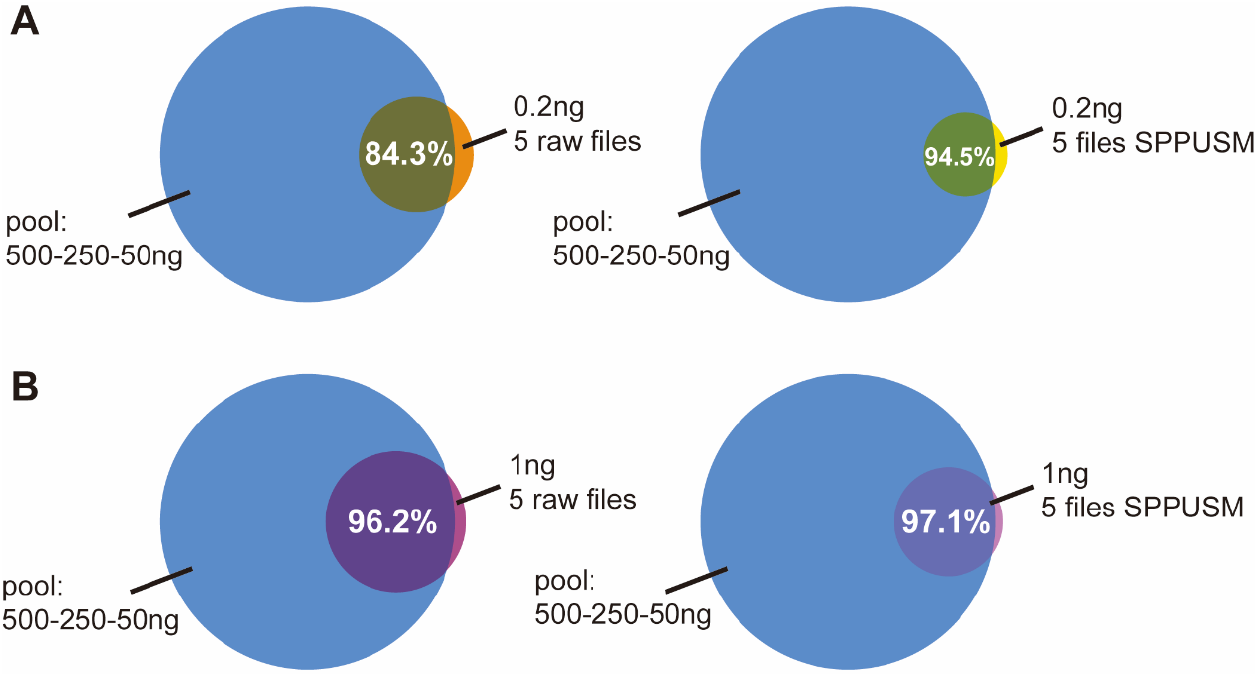
(A and B) Coverage of the peptides identified by SPPUSM and the peptide pool obtained using the classic database searching method.

After demonstrating the feasibility of using the SPPUSM workflow for improving spectral quality and peptide/protein identification, we further applied this method to single-cell proteome (SCP) analysis. In the SCP study, a large number of replicate tests on individual cells are conducted; therefore, plenty of data can be used for spectra merging without the need of extra sample analysis. Figure 4A exhibited the PSM, peptide and protein identification enhancement for each SCP file using SPPUSM. 28%-61% enhancements were achieved for protein identification for each SCP data. SCP data from other labs were also tested against the SPPUSM workflow to further support its capability. We used rare cell (50 HeLa cell) data from Zhu and Kelly’s lab^50^ (PXD016921), and a similar trend as that obtained in our own data was discovered. Although there was a lower extent of identification enhancement comparing with our own data, 12.0% improvement in protein identification was still achieved (Figure 4B). Lastly, we tested the ability of the SPPUSM workflow to detect the cellular heterogeneity of HeLa and HEK293T cells at the single-cell level. As shown in Figure 5, principal component analysis (PCA) using the newly identified proteins by the SPPUSM workflow resulted in the clear separation of the two types of cells, indicating the value of these lower abundant proteins for cellular functional investigation.

**Figure 4.**
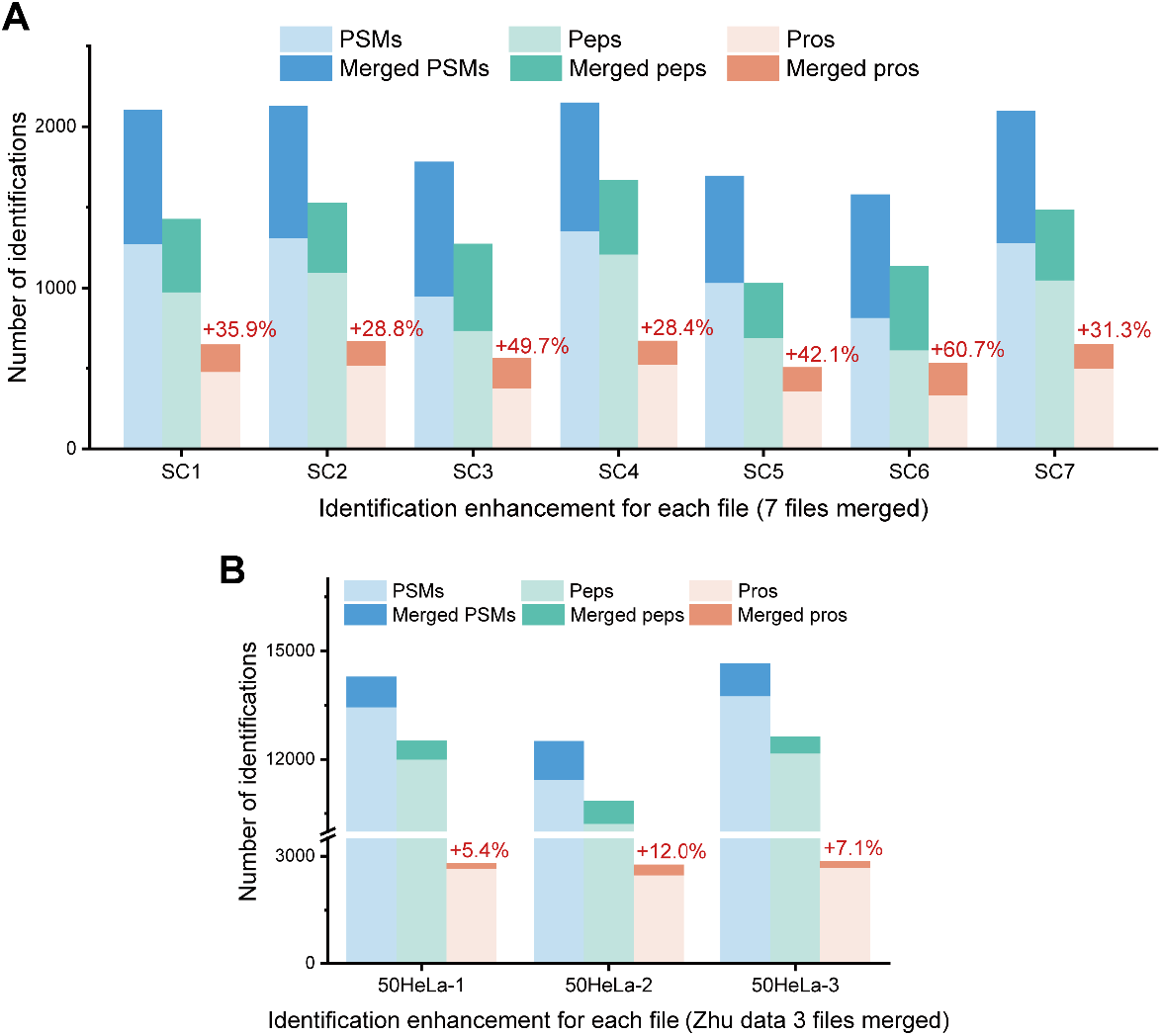
(A) PSM, peptide and protein identification enhancement by SPPUSM for each SCP data file obtained in this work. (B) Number of identifications obtained by the classic database searching strategy and by the SPPUSM workflow using the rare cell data from Zhu and Kelly’s lab.

**Figure 5.**
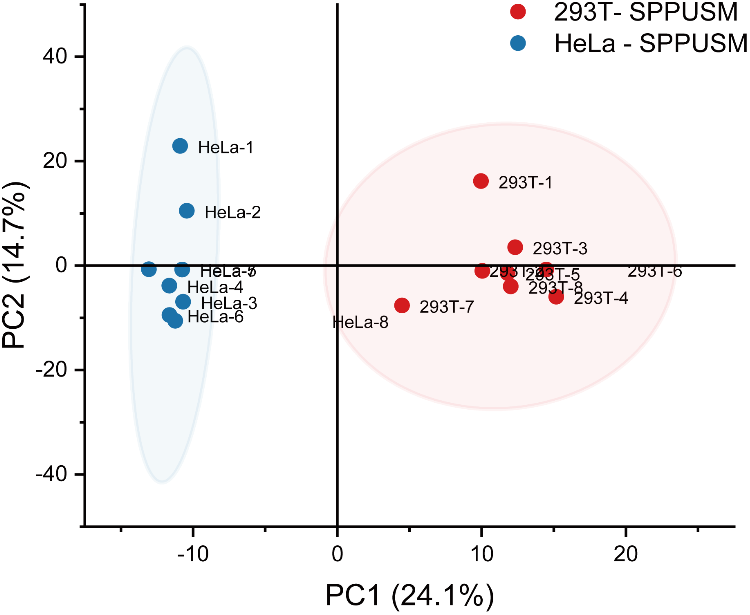
Cellular heterogeneity investigation of HeLa and HEK293T cells at the SCP level using the SPPUSM identified proteins.

## Conclusions

In this study, we developed the SPPUSM workflow as an MS data processing strategy for improving the proteome coverage of SCP and low-input samples. Applying this strategy led to 18.2% enhancements in protein identifications for 1 ng HeLa digests and the successful differentiation of cellular heterogeneity at the single-cell level. The combination of the SPPUSM workflow with advanced SCP sample processing and MS methods such as nanoPOTS and SCoPE2 should further enhance the sensitivity of SCP and promote its development.

## Acknowledgments

This study was supported by the National Natural Science Foundation of China (Nos. 32088101 and 22074158), the National Key R&D Program of China (Nos. 2021YFA1302604 and 2021YFA1301601) and the National Center for Protein Science (Beijing) Grant 2021-NCPSB-003. We thank Chunguang Han from the flow cytometry facility of the National Center for Protein Sciences Beijing for her help and technical support.

## Notes

The authors declare no competing financial interest.

The MS data have been deposited in the ProteomeXchange Consortium via the PRIDE partner repository under the dataset identifier PXD039336. All other data supporting the findings of this study are available from the corresponding author upon reasonable request.

## Graphical abstract

## Notes

### Competing Interest Statement

The authors have declared no competing interest.

## References

(1) Li, Y.; Xia, C.; Zhao, H.; Xie, Y.; Zhang, Y.; Zhang, W.; Yu, Y.; Wang, J.; Qin, W. A New Photolabeling Probe for Efficient Enrichment and Deep Profiling of Cell Surface Membrane Proteome by Mass Spectrometry. Chinese Chem. Lett. 2023, 34 (2), 107377. https://doi.org/10.1016/J.CCLET.2022.03.100.

(2) Dou, M.; Clair, G.; Tsai, C. F.; Xu, K.; Chrisler, W. B.; Sontag, R. L.; Zhao, R.; Moore, R. J.; Liu, T.; Pasa-Tolic, L.; Smith, R. D.; Shi, T.; Adkins, J. N.; Qian, W. J.; Kelly, R. T.; Ansong, C.; Zhu, Y. High-Throughput Single Cell Proteomics Enabled by Multiplex Isobaric Labeling in a Nanodroplet Sample Preparation Platform. Anal. Chem. 2019, 91 (20), 13119–13127. https://doi.org/10.1021/acs.analchem.9b03349.

(3) Specht, H.; Emmott, E.; Petelski, A. A.; Huffman, R. G.; Perlman, D. H.; Serra, M.; Kharchenko, P.; Koller, A.; Slavov, N. Single-Cell Proteomic and Transcriptomic Analysis of Macrophage Heterogeneity Using SCoPE2. Genome Biol. 2021, 22 (1), 1–27. https://doi.org/10.1186/s13059-021-02267-5.

(4) Wei, X.; Yang, M.; Jiang, Z.; Liu, J.; Zhang, X.; Chen, M.; Wang, J. A Modular Single-Cell Pipette Microfluidic Chip Coupling to ETAAS and ICP-MS for Single Cell Analysis. Chinese Chem. Lett. 2022, 33 (3), 1373–1376. https://doi.org/10.1016/J.CCLET.2021.08.024.

(5) Guo, Y.; Chen, P.; Gao, Z.; Li, Y.; Li, S.; Feng, X.; Liu, B. F. A Time-Coded Multi-Concentration Microfluidic Chemical Waveform Generator for High-Throughput Probing Suspension Single-Cell Signaling. Chinese Chem. Lett. 2022, 33 (6), 3091–3096. https://doi.org/10.1016/J.CCLET.2021.09.080.

(6) Zhou, W.; Lin, J.; Zhao, K.; Jin, K.; He, Q.; Hu, Y.; Feng, G.; Cai, Y.; Xia, C.; Liu, H.; Shen, W.; Hu, X.; Ouyang, H. Single-Cell Profiles and Clinically Useful Properties of Human Mesenchymal Stem Cells of Adipose and Bone Marrow Origin. Am. J. Sports Med. 2019, 47 (7), 1722–1733. https://doi.org/10.1177/0363546519848678.

(7) Aldridge, S.; Teichmann, S. A. Single Cell Transcriptomics Comes of Age. Nat. Commun. 2020, 11 (1). https://doi.org/10.1038/S41467-020-18158-5.

(8) Tanay, A.; Sebé-Pedrós, A. Evolutionary Cell Type Mapping with Single -Cell Genomics. Trends Genet. 2021, 37 (10), 919–932. https://doi.org/10.1016/J.TIG.2021.04.008.

(9) Hamey, F. K.; Lau, W. W. Y.; Kucinski, I.; Wang, X.; Diamanti, E.; Wilson, N. K.; Göttgens, B.; Dahlin, J. S. Single -Cell Molecular Profiling Provides a High-Resolution Map of Basophil and Mast Cell Development. Allergy 2021, 76 (6), 1731–1742. https://doi.org/10.1111/ALL.14633.

(10) Wang, C.; Hu, W.; Guan, L.; Yang, X.; Liang, Q. Single-Cell Metabolite Analysis on a Microfluidic Chip. Chinese Chem. Lett. 2022, 33 (6), 2883– 2892. https://doi.org/10.1016/J.CCLET.2021.10.006.

(11) Elowitz, M. B.; Levine, A. J.; Siggia, E. D.; Swain, P. S. Stochastic Gene Expression in a Single Cell. Science 2002, 297 (5584), 1183–1186. https://doi.org/10.1126/SCIENCE.1070919.

(12) Raj, A.; van Oudenaarden, A. Nature, Nurture, or Chance: Stochastic Gene Expression and Its Consequences. Cell 2008, 135 (2), 216–226. https://doi.org/10.1016/J.CELL.2008.09.050.

(13) Levy, E.; Slavov, N. Single Cell Protein Analysis for Systems Biology. Essays Biochem. 2018, 62 (4), 595–605. https://doi.org/10.1042/EBC20180014.

(14) Zhu, Y.; Piehowski, P. D.; Kelly, R. T.; Qian, W. J. Nanoproteomics Comes of Age. Expert Rev. Proteomics 2018, 15 (11), 865–871. https://doi.org/10.1080/14789450.2018.1537787.

(15) Budnik, B.; Levy, E.; Harmange, G.; Slavov, N. SCoPE-MS: Mass Spectrometry of Single Mammalian Cells Quantifies Proteome Heterogeneity during Cell Differentiation. Genome Biol. 2018, 19 (1), 1–12. https://doi.org/10.1186/S13059-018-1547-5/FIGURES/4.

(16) Zhang, P.; Gaffrey, M. J.; Zhu, Y.; Chrisler, W. B.; Fillmore, T. L.; Yi, L.; Nicora, C. D.; Zhang, T.; Wu, H.; Jacobs, J.; Tang, K.; Kagan, J.; Srivastava, S.; Rodland, K. D.; Qian, W. J.; Smith, R. D.; Liu, T.; Wiley, H. S.; Shi, T. Carrier-Assisted Single-Tube Processing Approach for Targeted Proteomics Analysis of Low Numbers of Mammalian Cells. Anal. Chem. 2019, 91 (2), 1441–1451. https://doi.org/10.1021/ACS.ANALCHEM.8B04258/SUPPL_FILE/AC8B04258_SI_002.PDF.

(17) Kulak, N. A.; Geyer, P. E.; Mann, M. Loss-Less Nano-Fractionator for High Sensitivity, High Coverage Proteomics* □ S Technological Innovation and Resources. Mol. Cell. Proteomics 2017, 16, 694–705. https://doi.org/10.1074/mcp.O116.065136.

(18) Martin, K.; Zhang, T.; Lin, T. T.; Habowski, A. N.; Zhao, R.; Tsai, C. F.; Chrisler, W. B.; Sontag, R. L.; Orton, D. J.; Lu, Y. J.; Rodland, K. D.; Yang, B.; Liu, T.; Smith, R. D.; Qian, W. J.; Waterman, M. L.; Wiley, H. S.; Shi, T. Facile One-Pot Nanoproteomics for Label-Free Proteome Profiling of 50-1000 Mammalian Cells. J. Proteome Res. 2021, 20 (9), 4452–4461. https://doi.org/10.1021/ACS.JPROTEOME.1C00403/SUPPL_FILE/PR1C00403_SI_004.XLSX.

(19) Specht, H.; Slavov, N. Transformative Opportunities for Single-Cell Proteomics. J. Proteome Res. 2018, 17 (8), 2565–2571. https://doi.org/10.1021/acs.jproteome.8b00257.

(20) Chen, Q.; Yan, G.; Gao, M.; Zhang, X. Ultrasensitive Proteome Profiling for 100 Living Cells by Direct Cell Injection, Online Digestion and Nano-LC-MS/MS Analysis. Anal. Chem. 2015, 87 (13), 6674–6680. https://doi.org/10.1021/ACS.ANALCHEM.5B00808/SUPPL_FILE/AC5B00808_SI_003.XLS.

(21) Zhu, Y.; Piehowski, P. D.; Zhao, R.; Chen, J.; Shen, Y.; Moore, R. J.; Shukla, K.; Petyuk, V. A.; Campbell-Thompson, M.; Mathews, C. E.; Smith, R. D.; Qian, W. J.; Kelly, R. T. Nanodroplet Processing Platform for Deep and Quantitative Proteome Profiling of 10–100 Mammalian Cells. Nat. Commun. 2018 91 2018, 9 (1), 1–10. https://doi.org/10.1038/s41467-018-03367-w.

(22) Li, Z. Y.; Huang, M.; Wang, X. K.; Zhu, Y.; Li, J. S.; Wong, C. C. L.; Fang, Q. Nanoliter-Scale Oil-Air-Droplet Chip-Based Single Cell Proteomic Analysis. Anal. Chem. 2018, 90 (8), 5430–5438. https://doi.org/10.1021/ACS.ANALCHEM.8B00661/SUPPL_FILE/AC8B00661_SI_001.PDF.

(23) Zy, L.; M, H.; Xk, W.; y, Z.; Js, L.; Ccl, W.; q, F. Nanoliter-Scale Oil-Air-Droplet Chip-Based Single Cell Proteomic Analysis. Anal. Chem. 2018, 90 (8), 5430–5438. https://doi.org/10.1021/ACS.ANALCHEM.8B00661.

(24) Michal Greguš, James C. Kostas, Somak Ray, Susan E. Abbatiello A. R. I. Improved Sensitivity of Ultralow Flow LC–MS-Based Proteomic Profiling of Limited Samples Using Monolithic Capillary Columns and FAIMS Technology. Anal. Chem. 2020, 92 (21), 14702–14712. https://doi.org/10.1021/acs.analchem.0c03262.Improved.

(25) Sun, B.; Kovatch, J. R.; Badiong, A.; Merbouh, N. Optimization and Modeling of Quadrupole Orbitrap Parameters for Sensitive Analysis toward Single-Cell Proteomics. J. Proteome Res. 2017, 16 (10), 3711–3721. https://doi.org/10.1021/acs.jproteome.7b00416.

(26) Swearingen, K. E.; Hoopmann, M. R.; Johnson, R. S.; Saleem, R. A.; Aitchison, J. D.; Moritz, R. L. Nanospray FAIMS Fractionation Provides Significant Increases in Proteome Coverage of Unfractionated Complex Protein Digests *. Mol. Cell. Proteomics 2012, 11 (4), M111.014985. https://doi.org/10.1074/MCP.M111.014985.

(27) Bonneil, E.; Pfammatter, S.; Thibault, P. Enhancement of Mass Spectrometry Performance for Proteomic Analyses Using High-Field Asymmetric Waveform Ion Mobility Spectrometry (FAIMS). J. Mass Spectrom. 2015, 50 (11), 1181– 1195. https://doi.org/10.1002/JMS.3646.

(28) Vasilopoulou, C. G.; Sulek, K.; Brunner, A.-D.; Meitei, N. S.; Schweiger-Hufnagel, U.; Meyer, S. W.; Barsch, A.; Mann, M.; Meier, F. Trapped Ion Mobility Spectrometry and PASEF Enable In-Depth Lipidomics from Minimal Sample Amounts. Nat. Commun. 2020 111 2020, 11 (1), 1–11. https://doi.org/10.1038/s41467-019-14044-x.

(29) Schoof, E. M.; Furtwängler, B.; Üresin, N.; Rapin, N.; Savickas, S.; Gentil, C.; Lechman, E.; Keller, U. auf dem; Dick, J. E.; Porse, B. T. Quantitative Single-Cell Proteomics as a Tool to Characterize Cellular Hierarchies. Nat. Commun. 2021, 12 (1), 1–15. https://doi.org/10.1038/s41467-021-23667-y.

(30) Siyal, A. A.; Chen, E. S. W.; Chan, H. J.; Kitata, R. B.; Yang, J. C.; Tu, H. L.; Chen, Y. J. Sample Size-Comparable Spectral Library Enhances Data-Independent Acquisition-Based Proteome Coverage of Low-Input Cells. Anal. Chem. 2021, 93 (51), 17003–17011. https://doi.org/10.1021/ACS.ANALCHEM.1C03477/ASSET/IMAGES/LARGE/AC1C03477_0007.JPEG.

(31) Watt, D. Van Der; Boekweg, H.; Truong, T.; Guise, A. J.; Plowey, E. D.; Kelly, R. T.; Payne, S. H. Benchmarking PSM Identification Tools for Single Cell Proteomics. bioRxiv 2021, 2021.08.17.456676. https://doi.org/10.1101/2021.08.17.456676.

(32) Boekweg, H.; Watt, D. Van Der; Truong, T.; Guise, A. J.; Plowey, E. D.; Kelly, R. T.; Payne, S. H. Features of Peptide Fragmentation Spectra in Single Cell Proteomics. bioRxiv 2021, 2021.08.17.456675. https://doi.org/10.1101/2021.08.17.456675.

(33) Williams, S. M.; Liyu, A. V.; Tsai, C. F.; Moore, R. J.; Orton, D. J.; Chrisler, W. B.; Gaffrey, M. J.; Liu, T.; Smith, R. D.; Kelly, R. T.; Pasa-Tolic, L.; Zhu, Y. Automated Coupling of Nanodroplet Sample Preparation with Liquid Chromatography-Mass Spectrometry for High-Throughput Single-Cell Proteomics. Anal. Chem. 2020, 92 (15), 10588–10596. https://doi.org/10.1021/ACS.ANALCHEM.0C01551/SUPPL_FILE/AC0C01551_SI_001.PDF.

(34) Lim, M. Y.; Paulo, J. A.; Gygi, S. P. Evaluating False Transfer Rates from the Match-between-Runs Algorithm with a Two-Proteome Model. J. Proteome Res. 2019, 18 (11), 4020–4026. https://doi.org/10.1021/ACS.JPROTEOME.9B00492/SUPPL_FILE/PR9B00492_SI_002.PDF.

(35) Bittremieux, W.; May, D. H.; Bilmes, J.; Noble, W. S. A Learned Embedding for Efficient Joint Analysis of Millions of Mass Spectra. Nat. Methods 2022 196 2022, 19 (6), 675–678. https://doi.org/10.1038/s41592-022-01496-1.

(36) Griss, J.; Perez-Riverol, Y.; Lewis, S.; Tabb, D. L.; Dianes, J. A.; Del-Toro, N.; Rurik, M.; Walzer, M.; Kohlbacher, O.; Hermjakob, H.; Wang, R.; Vizcano, J. Recognizing Millions of Consistently Unidentified Spectra across Hundreds of Shotgun Proteomics Datasets. Nat. Methods 2016 138 2016, 13 (8), 651– 656. https://doi.org/10.1038/nmeth.3902.

(37) Frank, A. M.; Monroe, M. E.; Shah, A. R.; Carver, J. J.; Bandeira, N.; Moore, R. J.; Anderson, G. A.; Smith, R. D.; Pevzner, P. A. Spectral Archives: Extending Spectral Libraries to Analyze Both Identified and Unidentified Spectra. Nat. Methods 2011 87 2011, 8 (7), 587–591. https://doi.org/10.1038/nmeth.1609.

(38) MacCoss, M. J.; Wu, C. C.; Yates, J. R. Probability-Based Validation of Protein Identifications Using a Modified SEQUEST Algorithm. Anal. Chem. 2002, 74 (21), 5593–5599. https://doi.org/10.1021/AC025826T.

(39) Cox, J.; Neuhauser, N.; Michalski, A.; Scheltema, R. A.; Olsen, J. V.; Mann, M. Andromeda: A Peptide Search Engine Integrated into the MaxQuant Environment. J. Proteome Res. 2011, 10 (4), 1794–1805. https://doi.org/10.1021/pr101065j.

(40) Kong, A. T.; Leprevost, F. V.; Avtonomov, D. M.; Mellacheruvu, D.; Nesvizhskii, A. I. MSFragger: Ultrafast and Comprehensive Peptide Identification in Mass Spectrometry-Based Proteomics. Nat. Methods 2017, 14 (5), 513–520. https://doi.org/10.1038/NMETH.4256.

(41) Chi, H.; Liu, C.; Yang, H.; Zeng, W. F.; Wu, L.; Zhou, W. J.; Wang, R. M.; Niu, X. N.; Ding, Y. H.; Zhang, Y.; Wang, Z. W.; Chen, Z. L.; Sun, R. X.; Liu, T.; Tan, G. M.; Dong, M. Q.; Xu, P.; Zhang, P. H.; He, S. M. Comprehensive Identification of Peptides in Tandem Mass Spectra Using an Efficient Open Search Engine. Nat. Biotechnol. 2018, 36 (11), 1059–1066. https://doi.org/10.1038/nbt.4236.

(42) Ye, X.; Yang, Y.; Zhou, J.; Xu, L.; Wu, L.; Huang, P.; Feng, C.; Ke, P.; He, A.; Li, G.; Li, Y.; Li, Y.; Lam, H.; Zhang, X.; Tian, R. Combinatory Strategy Using Nanoscale Proteomics and Machine Learning for T Cell Subtyping in Peripheral Blood of Single Multiple Myeloma Patients. Anal. Chim. Acta 2021, 1173. https://doi.org/10.1016/j.aca.2021.338672.

(43) An, M.; Zou, X.; Wang, Q.; Zhao, X.; Wu, J.; Xu, L. M.; Shen, H. Y.; Xiao, X.; He, D.; Ji, J. High-Confidence de Novo Peptide Sequencing Using Positive Charge Derivatization and Tandem MS Spectra Merging. Anal. Chem. 2013, 85 (9), 4530–4537. https://doi.org/10.1021/AC4001699/SUPPL_FILE/AC4001699_SI_001.PDF.

(44) Yang, H.; Li, Y. C.; Zhao, M. Z.; Wu, F. L.; Wang, X.; Xiao, W. Di; Wang, Y. H.; Zhang, J. L.; Wang, F. Q.; Xu, F.; Zeng, W. F.; Overall, C. M.; He, S. M.; Chi, H.; Xu, P. Precision de Novo Peptide Sequencing Using Mirror Proteases of Ac-Lysarginase and Trypsin for Large-Scale Proteomics. Mol. Cell. Proteomics 2019, 18 (4), 773–785. https://doi.org/10.1074/mcp.TIR118.000918.

(45) Yan, Y.; Kusalik, A. J.; Wu, F. X. De Novo Peptide Sequencing Using CID and HCD Spectra Pairs. Proteomics 2016, 16 (20), 2615–2624. https://doi.org/10.1002/PMIC.201500251.

(46) Zhang, W.; Yang, C.; Liu, J.; Liang, Z.; Shan, Y.; Zhang, L.; Zhang, Y. Accurate Discrimination of Leucine and Isoleucine Residues by Combining Continuous Digestion with Multiple MS3 Spectra Integration in Protein Sequence. Talanta 2022, 249, 123666. https://doi.org/10.1016/J.TALANTA.2022.123666.

(47) Wiśniewski, J. R.; Zougman, A.; Nagaraj, N.; Mann, M. Universal Sample Preparation Method for Proteome Analysis. Nat. Methods 2009, 6 (5), 359–362. https://doi.org/10.1038/nmeth.1322.

(48) Leduc, A.; Huffman, R. G.; Cantlon, J.; Khan, S.; Slavov, N. Exploring Functional Protein Covariation across Single Cells Using NPOP. Genome Biol. 2022 231 2022, 23 (1), 1–31. https://doi.org/10.1186/S13059-022-02817-5.

(49) Yu, F.; Fang, H.; Xiao, K.; Liu, Y.; Xue, B.; Tian, Z. Mass Measurement Accuracy of the Orbitrap in Intact Proteome Analysis. Rapid Commun. Mass Spectrom. 2016, 30 (12), 1391–1397. https://doi.org/10.1002/RCM.7574.

(50) Cong, Y.; Liang, Y.; Motamedchaboki, K.; Huguet, R.; Truong, T.; Zhao, R.; Shen, Y.; Lopez-Ferrer, D.; Zhu, Y.; Kelly, R. T. Improved Single-Cell Proteome Coverage Using Narrow-Bore Packed NanoLC Columns and Ultrasensitive Mass Spectrometry. Anal. Chem. 2020, 92 (3), 2665–2671. https://doi.org/10.1021/ACS.ANALCHEM.9B04631/SUPPL_FILE/AC9B04631_SI_002.XLSX.

