## Supplementary material for "SPPUSM: An MS/MS spectra merging strategy for improved low-input and single-cell proteome identification": Figure S1

**Experimental Section**

Synthetic standard peptides experiment

Eight synthetic standard peptides were obtained from Synpeptide Co. (China). All peptides were dissolved in 0.1% formic acid (Sigma Aldrich, USA) in water. Peptides ASLINPGQEK, GYGSNFVVGER, MAFDLERPGVPVENR and GYLQPYTDEEEDALIHR were mixed to produce a solution of 10ng/µL per peptide and 1µL of this solution was injected for LC-MS analysis. Peptides ASGNLIPQEK, GYGVVFSNGER, MAFRPGVVDLPEENR and GYTLADEEEDPLIYQHR were mixed in the same way for LC-MS/MS analysis too.

Sequence of the synthetic standard peptides:

H-Ala-Leu-Ile-Asn-Pro-Gly-Gln-Glu-Lys-OH (ASLINPGQEK),

H-Ala-Ser-Gly-Asn-Leu-Ile-Pro-Gln-Glu-Lys-OH (ASGNLIPQEK),

H-Gly-Tyr-Gly-Ser-Asn-Phe-Val-Val-Gly-Glu-Arg-OH (GYGSNFVVGER),

H-Gly-Tyr-Gly-Val-Val-Phe-Ser-Asn-Gly-Glu-Arg-OH (GYGVVFSNGER),

H-Met-Ala-Phe-Asp-Leu-Glu-Arg-Pro-Gly-Val-Pro-Val-Glu-Asn-Arg-OH (MAFDLERPGVPVENR),

H-Met-Ala-Phe-Arg-Pro-Gly-Val-Val-Asp-Leu-Pro-Glu-Glu-Asn-Arg-OH (MAFRPGVVDLPEENR),

H-Gly-Tyr-Leu-Gln-Pro-Tyr-Thr-Asp-Glu-Glu-Glu-Asp-Ala-Leu-Ile-His-Arg-OH (GYLQPYTDEEEDALIHR)

H-Gly-Tyr-Thr-Leu-Ala-Asp-Glu-Glu-Glu-Asp-Pro-Leu-Ile-Tyr-Gln-His-Arg-OH (GYTLADEEEDPLIYQHR),

**Supplementary Figures**

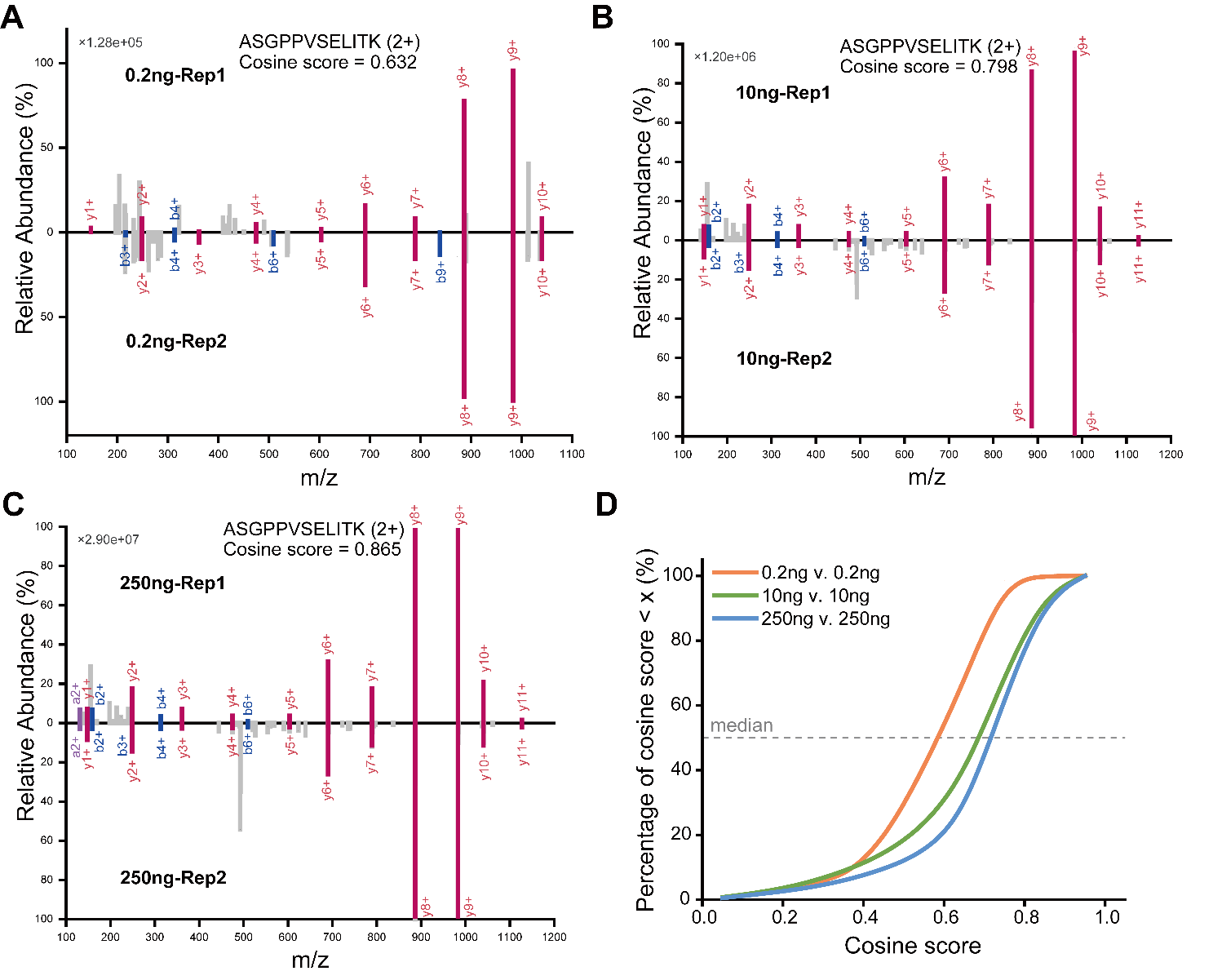

Figure S1. (A-C) Spectra patterns of one co-identified peptide in 0.2 ng, 10ng and 250 ng Hela digest sample. (D) Cosine scores of all identified PSMs from 0.2 ng, 10 ng and 250ng HeLa digest sample.

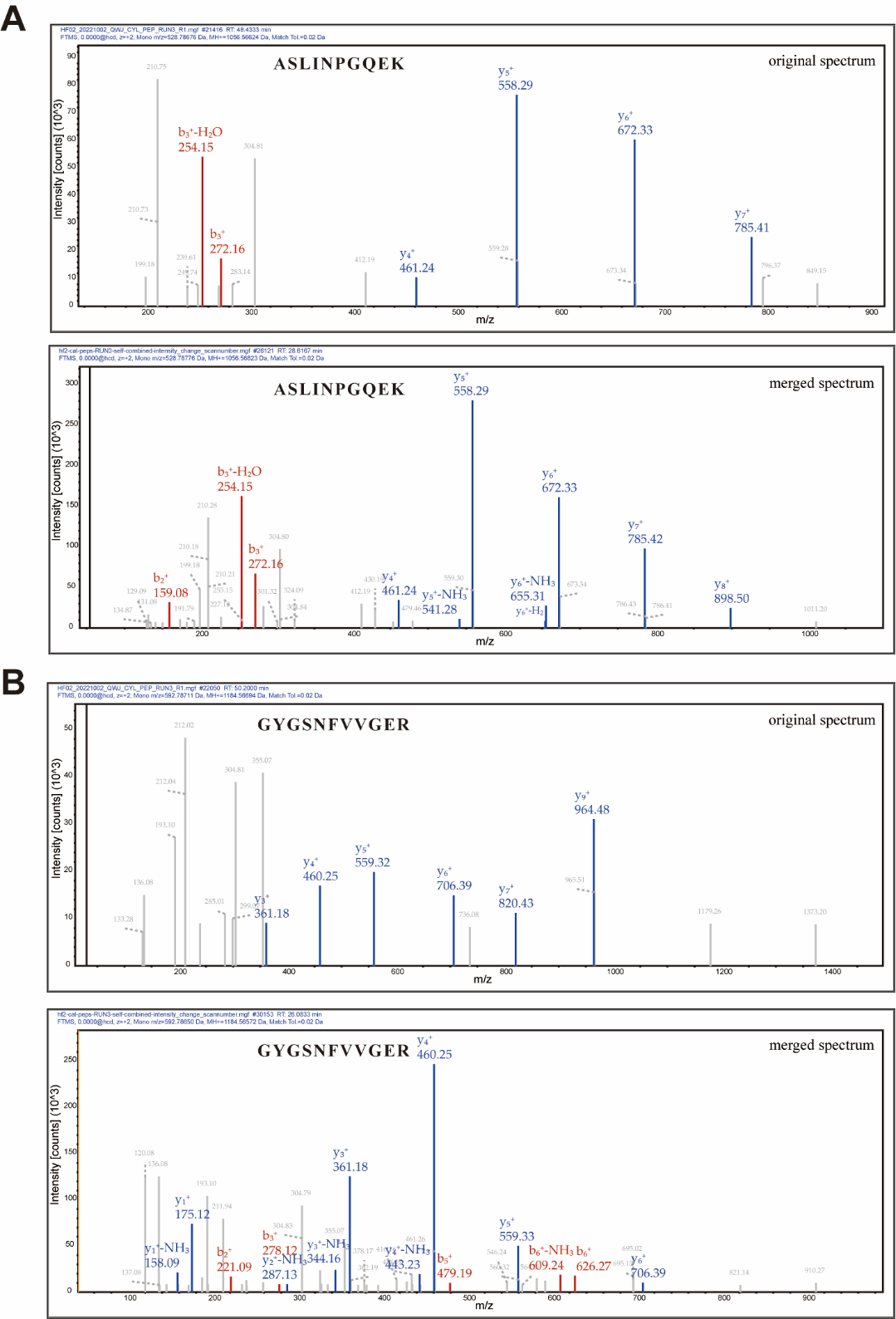

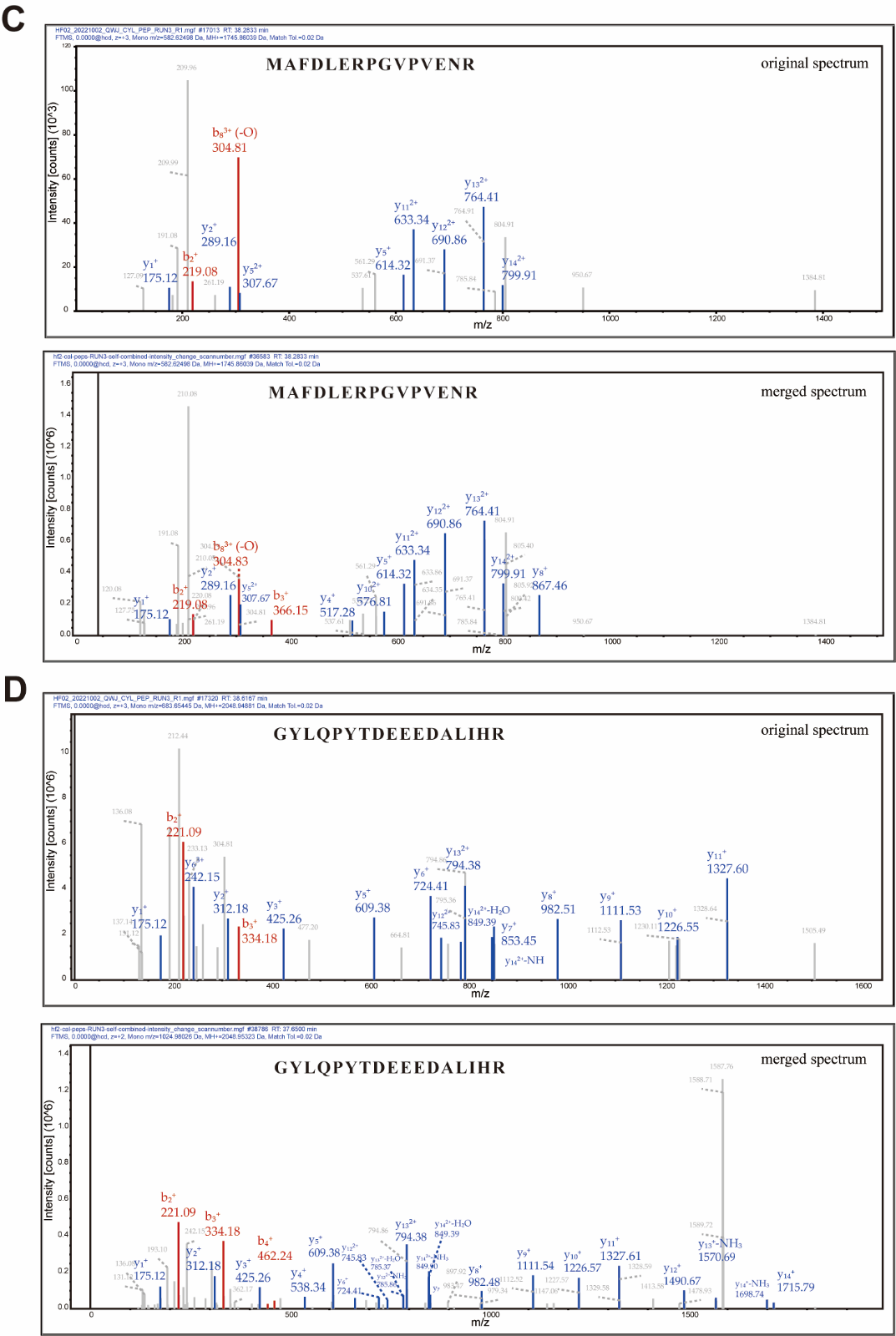

Figure S2. Identified MS/MS spectra for synthetic standard peptides before and after merging. (A) Peptide ASLINPGQEK. (B) Peptide GYGSNFVVGER. (C) Peptide MAFDLERPGVPVENR. (D) Peptide GYLQPYTDEEEDALIHR.

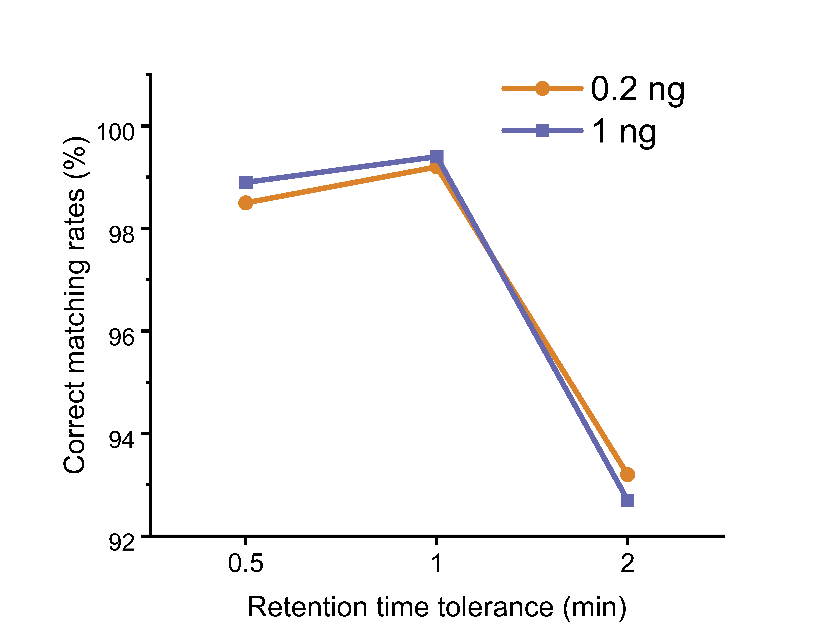

Figure S3. Correct matching rates using 2, 1 and 0.5 min RT tolerance.

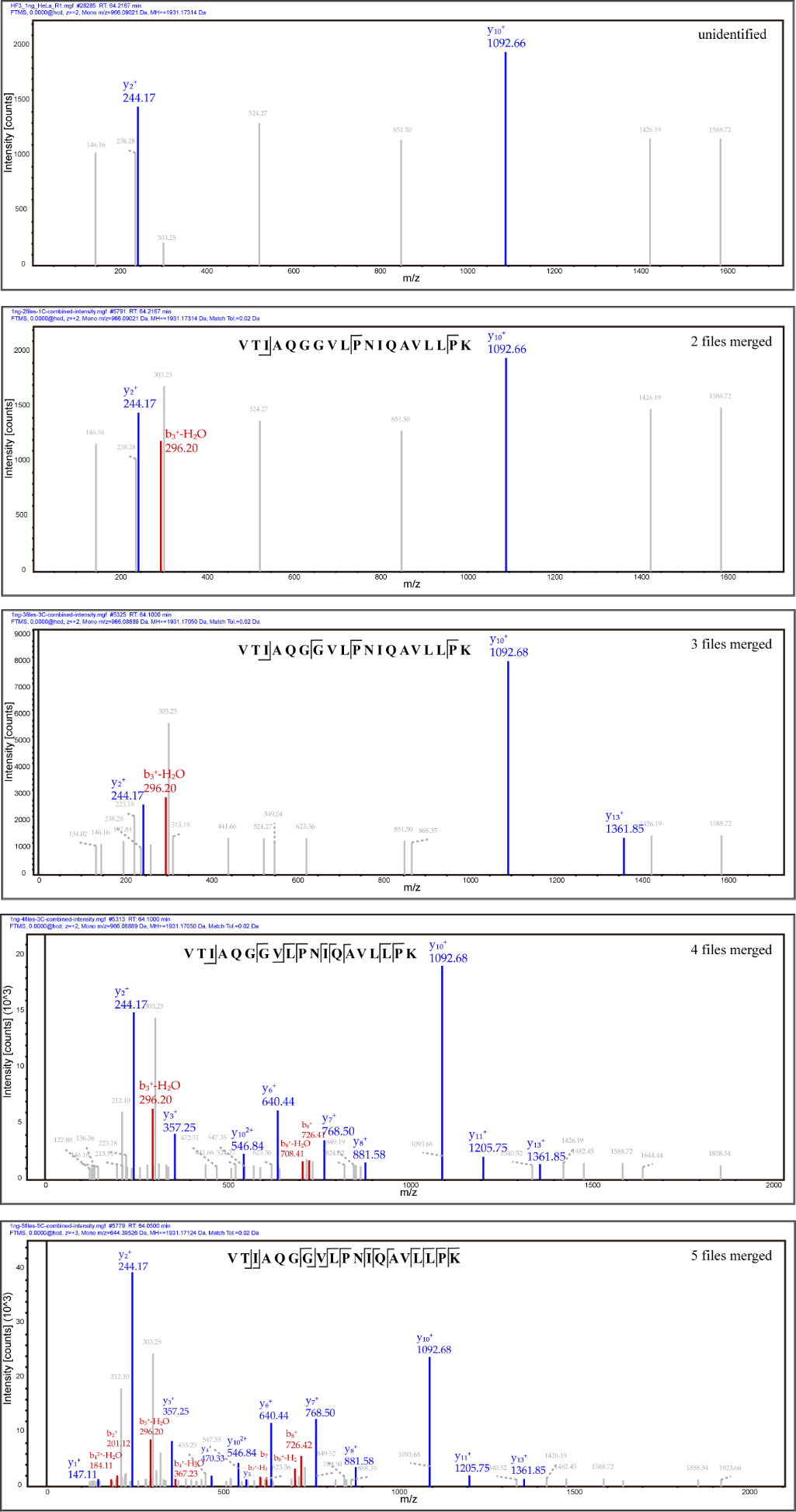

Figure S4. Typical MS/MS spectra of peptide VTIAQGGVLPNIQAVLLPK in 1 ng samples using different number of merging files.

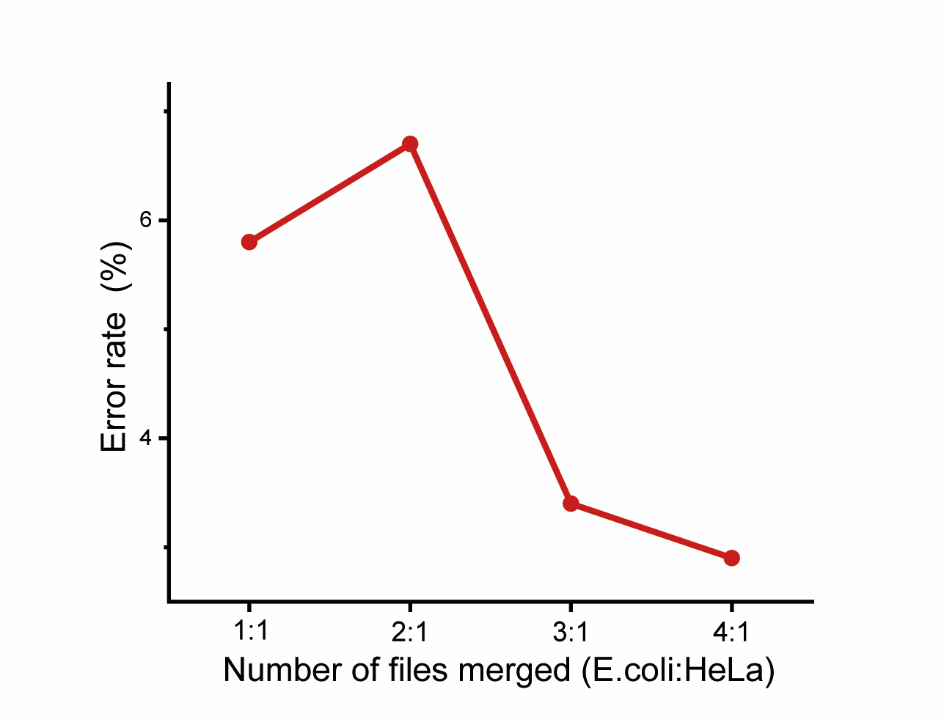

Figure S5. Error rates calculated using the peptides identified by the SPPUSM workflow using merged spectra from HeLa and E. coli digests.
